# NAChRDB: A Web Resource of Structure-Function Annotations to Unravel the Allostery of Nicotinic Acetylcholine Receptors

**DOI:** 10.1101/2020.01.08.898171

**Authors:** Aliaksei Chareshneu, Purbaj Pant, Ravi José Tristão Ramos, Tuğrul Gökbel, Crina-Maria Ionescu, Jaroslav Koča

## Abstract

**Summary:** Due to their paramount importance, near-ubiquitous presence, and complex nature, nicotinic acetylcholine receptors (nAChRs) have remained the focus of intensive research for over 50 years. The vast amount of knowledge accumulated on the topic has become extremely difficult to navigate. NAChRDB addresses this challenge by providing web-based, real-time access to curated residue-level functional annotations of neuromuscular nAChRs with interactive 3D visualization and sequence alignment. NAChRDB provides systematic access to experimental observations and predictions from computational studies reported in the literature or performed specifically to complement current knowledge, which allows new findings to be interpreted in a more holistic context, both from a structural and a functional perspective. NAChRDB aims to serve as an invaluable resource for identifying gaps in knowledge and for guiding discovery through structural and molecular biology experiments, especially when exploring the allosteric mechanisms underlying neuromuscular nAChR function and pathology.

**Availability and Implementation:** NAChRDB is freely available online, with a self-explanatory interface and useful tool tips (https://crocodile.ncbr.muni.cz/Apps/NAChRDB/). No installation or user registration is required. NAChRDB content is stored in .json format, queried using Python, and rendered in browser using Javascript and WebGL (LiteMol). NAChRDB is highly responsive and accessible through any modern Internet browser on desktop and mobile devices.

**Supplementary information:** Supplementary data are available online.

## 1 Introduction

The nicotinic acetylcholine receptor (nAChR) is an evolutionarily ancient allosteric membrane protein mediating synaptic transmission (Changeux, 2018). This prototypic member of the pentameric ligand-gated ion-channel family is involved in many physiological and pathological processes including neurological diseases and addictions. NAChRs convert the chemical signal of acetylcholine into ion current by allowing sodium and calcium ions to enter the cell and potassium ions to exit the cell. This fast conversion of chemical signal to ion current relies heavily on allosteric regulation, which links the agonist-binding pocket in the extracellular domain to a gating mechanism lying approximately 60 Å away in a central channel spanning the entire length of the molecule. Hundreds of nAChR-binding compounds have been found to modulate nerve impulse transmission by regulating this coupling of agonist binding and channel gating (Gündisch and Eibl, 2011; Reyes-Parada and Iturriaga-Vasquez, 2016). Not surprisingly, nAChRs have remained the focus of intensive research for more than 50 years (Brown, 2019). The effort to understand structure-function relationships in nAChRs has resulted in huge amounts of structural and functional information. However, the cumulative knowledge on nAChRs is not systematically accessible, with thousands upon thousands of experimental and/or computational studies reporting data on different receptor types, using different residue numbering schemes, and different terminology. Furthermore, because nAChRs are very large and difficult to crystalize, structural studies have focused on individual parts of the molecule, which makes it challenging to compile the current knowledge in an efficient manner and to promote further discoveries (Changeux, 2012; Halliwell, 2007). Finally, in the absence of comprehensive and unified structural annotation, it is almost impossible to identify gaps in knowledge or areas of controversy. We developed NAChRDB to address such limitations by providing web-based, real-time access to curated residue-level functional annotations of neuromuscular nAChRs, with interactive 3D visualization and sequence alignment.

## 2 Data Sources and Database Coverage

Functional annotations were collected from a semi-systematic literature review of Medline through PubMed (**see Supplementary Materials**). Upon manually scanning over 3000 papers on neuromuscular nAChRs, we selected 117 studies published between 1982 and 2019 and providing functional annotations of specific residues or parts of the protein. Currently, NAChRDB contains approximately 2000 unique annotation records describing the role of specific residues, as inferred based on experimental observations or computational predictions reported for nAChRs of organisms from 6 genera. These studies, which cover 41 experimental or computational methods, were conducted at 92 institutions from 25 countries. Additionally, we conducted two computational studies to complement current knowledge, which resulted in the addition of 741 annotation records. NAChRDB thus provides a comprehensive view on structure-function relationships in the neuromuscular nAChR (Fig. 1).

**Fig. 1.**
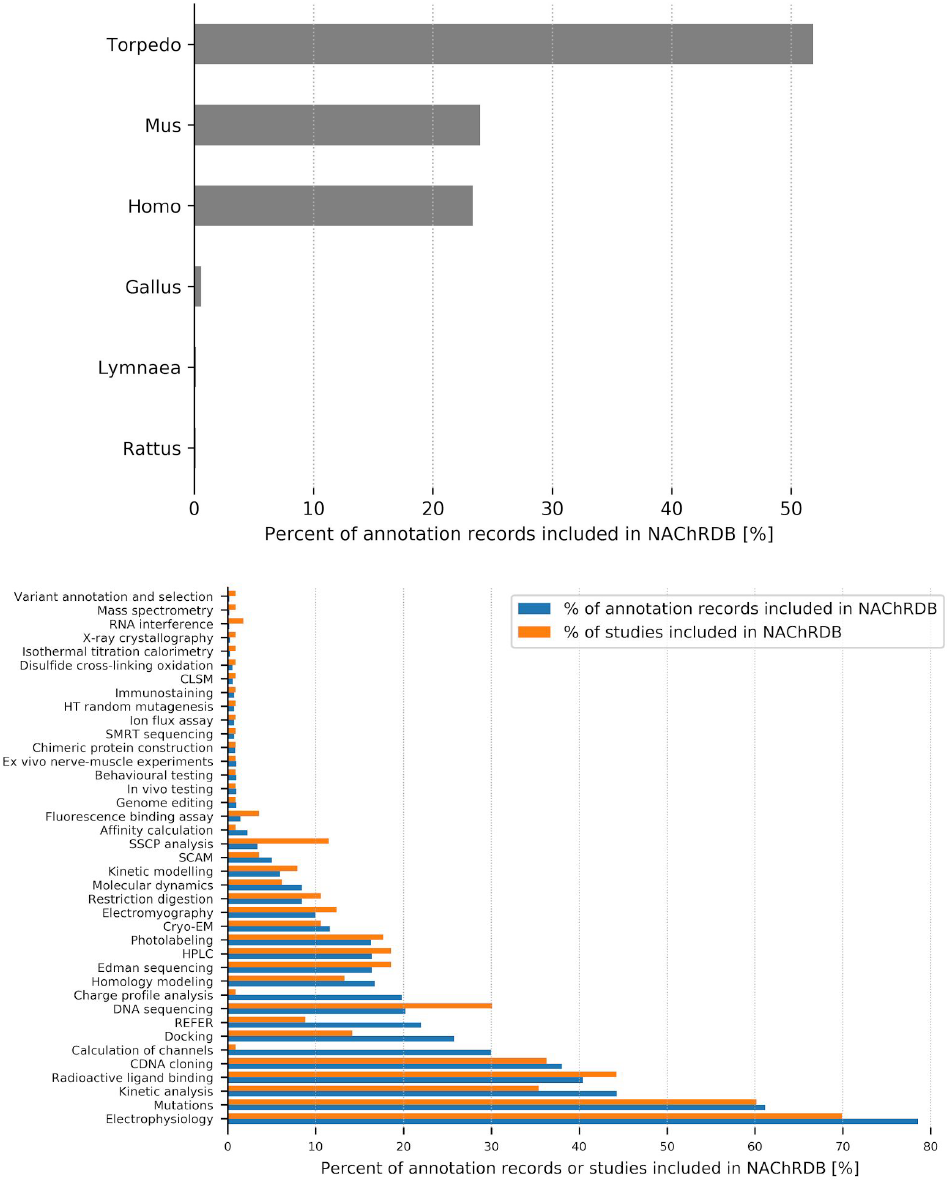
Current coverage of NAChRDB. NAChRDB provides an overview of residue-level structure-function annotations in neuromuscular nAChRs. *(Top)* Most annotation records were obtained from studies on the *Mus*, *Torpedo*, or *Homo* nAChRs *(Bottom)*. Whereas experiments involving electrophysiological measurements, mutagenesis, or radioactive ligands account for most studies covered by NAChRDB, many residue-level annotations were also obtained from studies based on less commonly used methods, such as docking or rate-equilibrium free energy relationship.

The structures and sequences of relevant nAChRs were obtained from the Protein Data Bank and from Uniprot, respectively. Sequence alignments were performed using Clustal W (Thompson et al., 1994). Residues potentially involved in charge transfer networks facilitating channel gating were predicted using a modified charge profile analysis (Ionescu et al., 2012). Briefly, the following steps were employed: (i) conformers were generated based on normal mode analysis using elNemo (Suhre and Sanejouand, 2004); atomic charges for each conformer were computed using AtomicChargeCalculator (Ionescu *et al.*, 2015) with several different settings; residue charges were then compared across conformers; outliers were reported based on robust statistical analysis (**see Supplementary Materials**). Channel lining residues were predicted using ChannelsDB (Pravda et al., 2018) (**see Supplementary Materials**). All predictions were added to NAChRDB as annotation records, with full reference to the source of information. In-house scripts were employed for data processing, but all annotations were curated manually.

## 3 Implementation

NAChRDB content is stored in. json format, queried using Python, and rendered in browser using Javascript and LiteMol, a WebGL-based technology for real-time in-browser rendering of large-scale macromolecular structures (Sehnal *et al.*, 2017) (Fig. 2). NAChRDB is freely available online (https://crocodile.ncbr.muni.cz/Apps/NAChRDB/), with no requirement for installation or user registration, with self-explanatory interface and useful tool tips. NAChRDB is highly responsive and accessible through any modern Internet browser on desktop and mobile devices.

**Fig. 2.**
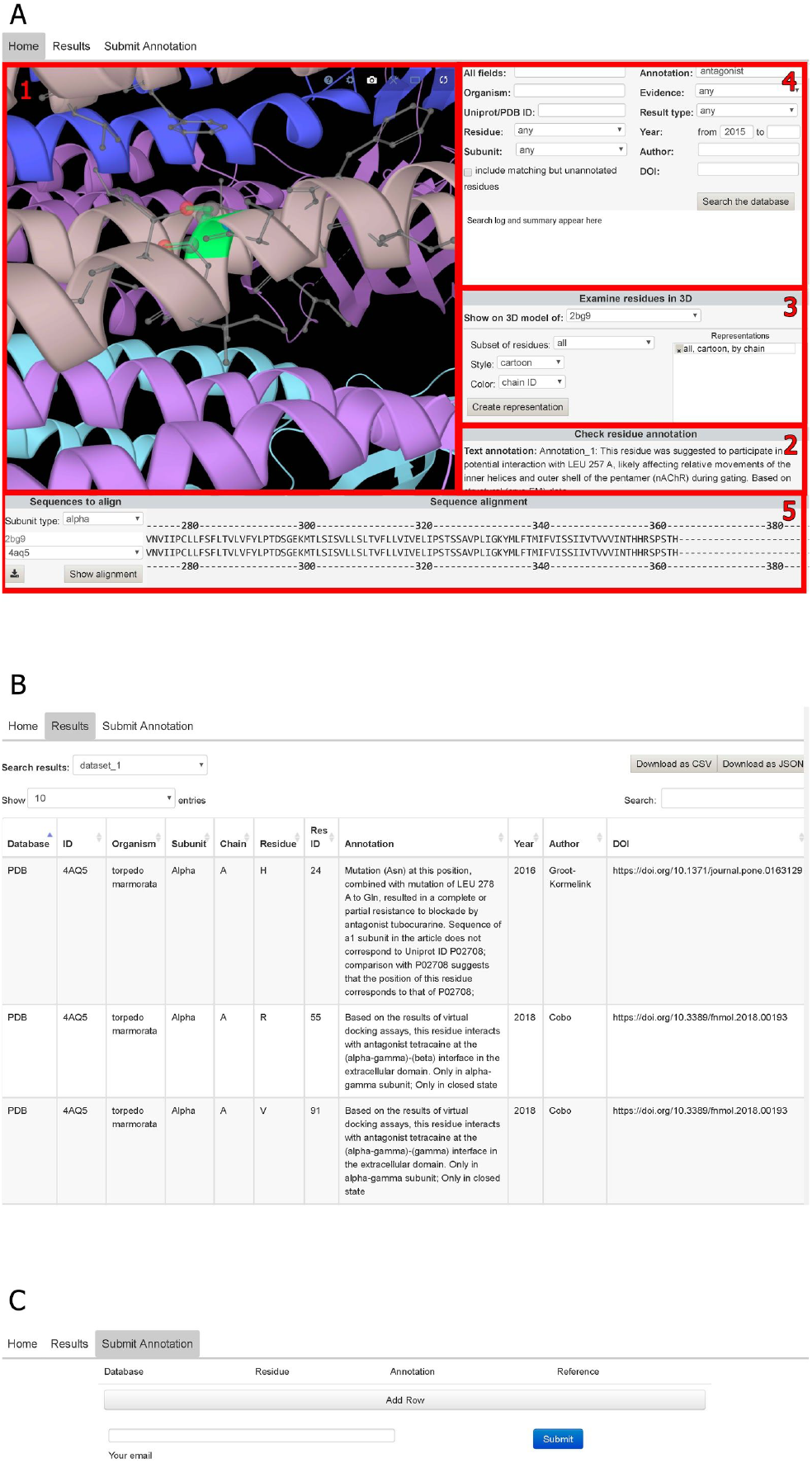
Screenshot of NAChRDB on mobile devices. The NAChRDB workspace is organized in several tabs. The Home tab *(A)* contains: (1) a 3D viewer widget providing interactive visualization of nAChR 3D models, complete with (2) direct reporting of annotation records for a selected residue and (3) 3D representation settings; (4) a search section providing extensive, PubMed-like search functionality for querying structure, function, and literature-related fields; (5) a sequence alignment viewer that also enables direct reporting of annotation records available for a selected residue. The Results tab *(B)* summarizes the search results in an interactive table. All results can be downloaded in. csv and. json format. The Submit tab *(C)* allows users to report annotations, thus contributing to maintaining NAChRDB up to date.

## 4 Case studies

1. A simple text search in NAChRDB helps to quickly gauge the potential role of specific residues, as well as to identify opportunities for further study. For example, certain residues found to link different domains, bind certain drug molecules, contribute to channel gating, form N-glycosylation sites, or regulate agonist binding were never studied in human neuromuscular nAChR (Fig. 3A).
2. When conducting new computational studies, NAChRDB helps to put the computational predictions into context, as well as to identify gaps in current knowledge and thus opportunities for further study. For example, among the residues predicted to line the channel’s inner surface in neuromuscular Torpedo nAChR, 28.5% have been studied to date; among the residues predicted to contribute to protein-wide charge transfer networks in Torpedo nAChR that facilitate channel gating, 18.5% have been studied (Fig. 3B).
3. Additionally, NAChRDB helps to identify potential areas of ambiguity or controversy in the current knowledge. For example, much remains to be clarified regarding the relationship between channel opening kinetics and voltage dependency, especially in nAChRs where key ASP residues are mutated to non-charged residues (Fig. 3C).

**Fig. 3.**
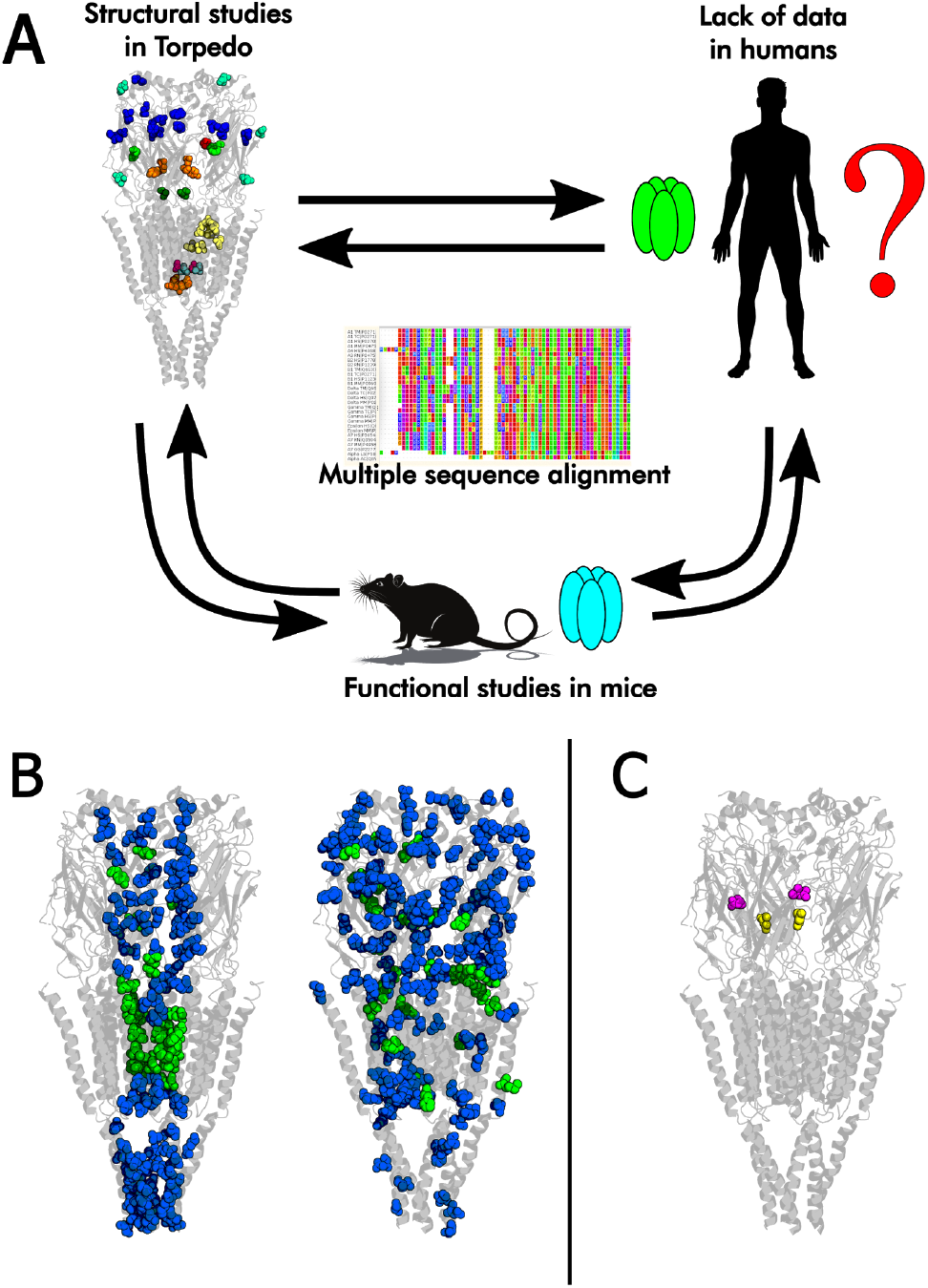
Case studies illustrating the usefulness of NAChRDB as a unified resource for structure-function annotations of neuromuscular nAChRs. (*A)* It is very easy to identify gaps in knowledge and design future investigations across different species. *Color coding on the 3D model:* dark green – αγVAL46 and αδVAL46 were suggested to serve as a key structural link between the extracellular and transmembrane domains of the neuromuscular nAChR in *Torpedo marmorata* (Miyazawa et al., 2003); teal and pink – αγSER248, αδSER248, βSER254, and δSER262 form a binding site for a channel-blocking non-competitive antagonist of the neuromuscular nAChR in *T. marmorata* (Hucho et al., 1986); orange – αγTYR127, αδTYR127, αγASP97, αδASP97, αγTHR244, αδTHR244, αγLYS242, and αδLYS242 participate in the channel gating process of neuromuscular nAChR in murines (Purohit and Auerbach, 2007; Zhang and Karlin, 1998); green – mutations of γTRP55 and δTRP57 change the agonist affinity of neuromuscular nAChR in murines (Nayak et al., 2016); red – mutations of δPRO123 disable one ACh binding site in murine neuromuscular nAChR (Gupta et al., 2013); cyan – γASN68, γASN141, δASN70, δASN143, and δASN208 may form N-glycosylation sites in the neuromuscular nAChR of *T. californica* (Chiara and Cohen, 1997; Strecker et al., 1994); blue – αγASP195, αδASP195, γLEU77, αγPRO194, αδPRO194, αγCYS193, αδCYS193, γLYS10, αγTRP86, αδTRP86, αγPRO88, αδPRO88, γTYR104, γILE80, and δTYR212 may contribute to physostigmine binding sites in the neuromuscular nAChR of *T. californica* (Hamouda et al., 2013); pale yellow and pink - δTHR274, δPHE232, δCYS236, δARG277, αγSER248, αδSER248, βSER254, βVAL261, and δVAL269 may form propofol binding sites in the neuromuscular nAChR of *T. californica* (Jayakar et al., 2013). The residue labels include the subunit(s), residue name, and residue number. Residue numbering is given according to the neuromuscular nAChR sequence from the respective organism (**see Supplementary Materials**). *(B)* Upon conducting computational studies to identify channel lining residues (left) and charge transfer networks (right), one can easily examine the predictions in the context of current knowledge (green), as well as identify areas of further study (blue). *(C)* Mutation of αASP200 (magenta) to GLN in the mouse neuromuscular nAChR expressed in human kidney cells was reported to have a dramatic effect on channel opening rate (Akk et al., 1996) but a non-significant effect on slope conductance or voltage dependency (Dworakowska et al., 2018). Some mutations of αASP97 (yellow) in mouse neuromuscular nAChR were shown to dramatically change gating kinetics (Chakrapani and Auerbach, 2005; Purohit and Auerbach, 2007), whereas other mutations were found to have only a marginal effect (Chakrapani et al., 2003). The 3D model of *T. marmorata* nAChR (PDB ID: 4AQ9) (Unwin and Fujiyoshi, 2012) was used for graphical representation in all panels.

Armed with such information, it is easy to identify understudied areas and design further experiments that can cover the current gaps and provide a more meaningful contribution to the current knowledge. Many residues highlighted by state-of-the-art computational analyses have not been investigated to date, mainly because experimental studies are expensive and thus typically focus only on areas thought to be of utmost importance. Furthermore, due to reporting bias, negative results are scarce. Thus, in addition to serving as an easy reference for structure-function information, we hope that NAChRDB will help promote the reporting of both positive and negative results, so that the scientific community may form a comprehensive picture of the functioning of nAChRs.

## 5 Limitations

At present, the annotation records in NAChRDB refer mainly to the neuromuscular receptor. Extending NAChRDB to cover neuronal receptors is our current focus. Moreover, users can submit suggestions for new annotations directly via NAChRDB; we will review these suggestions manually and update NAChRDB accordingly. We are also working on a text mining tool that will enable us to automatically expand the coverage of NAChRDB as soon as new information becomes available.

## 6 Conclusions

NAChRDB helps to quickly summarize the knowledge about specific parts of neuromuscular nAChRs, as well as to detect conflicting results reported for the same or homologous residues, and even to identify the parts of the protein not studied to date. NAChRDB provides systematic access to experimental observations and predictions from computational studies reported in the literature or performed specifically to complement current knowledge. New findings can thus be interpreted in a more holistic context, both from a structural and a functional perspective. Ultimately, NAChRDB aims to serve as an invaluable resource for guiding discovery through structural and molecular biology experiments, especially when exploring the allosteric mechanisms underlying nAChR function and pathology. In addition, NAChRDB can serve as a key starting point for the unification of the state-of-art knowledge in the broad field of pentameric ligand-gated ion channels.

## Supporting information

S1.pdb

S2.pdb

S3.csv

S4.csv

S5.csv

## Acknowledgements

The authors would like to thank Dr. David Sehnal for expert technical assistance in implementing the 3D viewer widget which is based on the LiteMol suite.

## References

Akk,G. et al. (1996) Binding sites contribute unequally to the gating of mouse nicotinic alpha D200N acetylcholine receptors. J. Physiol., 496 (Pt 1), 185–96.

Brown,D.A. (2019) Acetylcholine and cholinergic receptors. Brain Neurosci. Adv., https://doi.org/10.1177/2398212818820506

Chakrapani,S. et al. (2003) The role of loop 5 in acetylcholine receptor channel gating. J. Gen. Physiol., 122, 521–539.

Chakrapani,S. and Auerbach,A. (2005) A speed limit for conformational change of an allosteric membrane protein. Proc. Natl. Acad. Sci., 102, 87–92.

Changeux,J.-P. (2018) The nicotinic acetylcholine receptor: a typical ‘allosteric machine’. Philos. Trans. R. Soc. Lond. B. Biol. Sci., 373.

Changeux,J.-P. (2012) The nicotinic acetylcholine receptor: the founding father of the pentameric ligand-gated ion channel superfamily. J. Biol. Chem., 287, 40207–15.

Chiara,D.C. and Cohen,J.B. (1997) Identification of amino acids contributing to high and low affinity d-tubocurarine sites in the *Torpedo* nicotinic acetylcholine receptor. J. Biol. Chem., 272, 32940–50.

Domville,J.A. and Baenziger,J.E. (2018) An allosteric link connecting the lipid-protein interface to the gating of the nicotinic acetylcholine receptor. Sci. Rep., 8, 3898.

Dworakowska,B. et al. (2018) Hydrocortisone inhibition of wild-type and αD200Q nicotinic acetylcholine receptors. Chem. Biol. Drug Des., 92, 1610–1617.

Groot-Kormelink,P.J. et al. (2016) High throughput random mutagenesis and single molecule real time sequencing of the muscle nicotinic acetylcholine receptor. PLoS One, 11, e0163129.

Gündisch,D. and Eibl,C. (2011) Nicotinic acetylcholine receptor ligands, a patent review (2006-2011). Expert Opin. Ther. Pat., 21, 1867–96.

Gupta,S. et al. (2013) Function of interfacial prolines at the transmitter-binding sites of the neuromuscular acetylcholine receptor. J. Biol. Chem., 288, 12667–79.

Halliwell,R.F. (2007) A short history of the rise of the molecular pharmacology of ionotropic drug receptors. Trends Pharmacol. Sci., 28, 214–9.

Hamouda,A.K. et al. (2013) Physostigmine and galanthamine bind in the presence of agonist at the canonical and noncanonical subunit interfaces of a nicotinic acetylcholine receptor. J. Neurosci., 33, 485–494.

Hucho,F. et al. (1986) The ion channel of the nicotinic acetylcholine receptor is formed by the homologous helices M II of the receptor subunits. FEBS Lett., 205, 137–142.

Ionescu,C.-M. et al. (2015) AtomicChargeCalculator: interactive web-based calculation of atomic charges in large biomolecular complexes and drug-like molecules. J. Cheminform., 7, 50.

Ionescu,C.-M. et al. (2012) Charge profile analysis reveals that activation of pro-apoptotic regulators Bax and Bak relies on charge transfer mediated allosteric regulation. PLoS Comput. Biol., 8, e1002565.

Jayakar,S.S. et al. (2013) Identification of propofol binding sites in a nicotinic acetylcholine receptor with a photoreactive propofol analog. J. Biol. Chem., 288, 6178–6189.

Miyazawa,A. et al. (2003) Structure and gating mechanism of the acetylcholine receptor pore. Nature, 423, 949–955.

Nayak,T.K. et al. (2016) Structural correlates of affinity in fetal versus adult endplate nicotinic receptors. Nat. Commun., 7.

Pravda,L. et al. (2018) ChannelsDB: database of biomacromolecular tunnels and pores. Nucleic Acids Res., 46, D399–D405.

Purohit,P. and Auerbach,A. (2007) Acetylcholine receptor gating: Movement in the α-subunit extracellular domain. J. Gen. Physiol., 130, 569–579.

Reyes-Parada,M. and Iturriaga-Vasquez,P. (2016) The development of novel polypharmacological agents targeting the multiple binding sites of nicotinic acetylcholine receptors. Expert Opin. Drug Discov., 11, 969–81.

Sehnal,D. et al. (2017) LiteMol suite: interactive web-based visualization of large-scale macromolecular structure data. Nat. Methods, 14, 1121–1122.

Strecker,A. et al. (1994) All potential glycosylation sites of the nicotinic acetylcholine receptor δ subunit from *Torpedo californica* are utilized. Eur. J. Biochem., 220, 1005–1011.

Suhre,K. and Sanejouand,Y.-H. (2004) ElNemo: a normal mode web server for protein movement analysis and the generation of templates for molecular replacement. Nucleic Acids Res., 32, W610–4.

Thompson,J.D. et al. (1994) CLUSTAL W: Improving the sensitivity of progressive multiple sequence alignment through sequence weighting, position-specific gap penalties and weight matrix choice. Nucleic Acids Res., 22, 4673–4680.

Unwin,N. and Fujiyoshi,Y. (2012) Gating movement of acetylcholine receptor caught by plunge-freezing. J. Mol. Biol., 422, 617–634.

Zhang,H. and Karlin,A. (1998) Contribution of the beta subunit M2 segment to the ion-conducting pathway of the acetylcholine receptor. Biochemistry, 37, 7952–64.

