## Supplementary Materials for "NAChRDB: A Web Resource of Structure-Function Annotations to Unravel the Allostery of Nicotinic Acetylcholine Receptors"

### Literature scanning

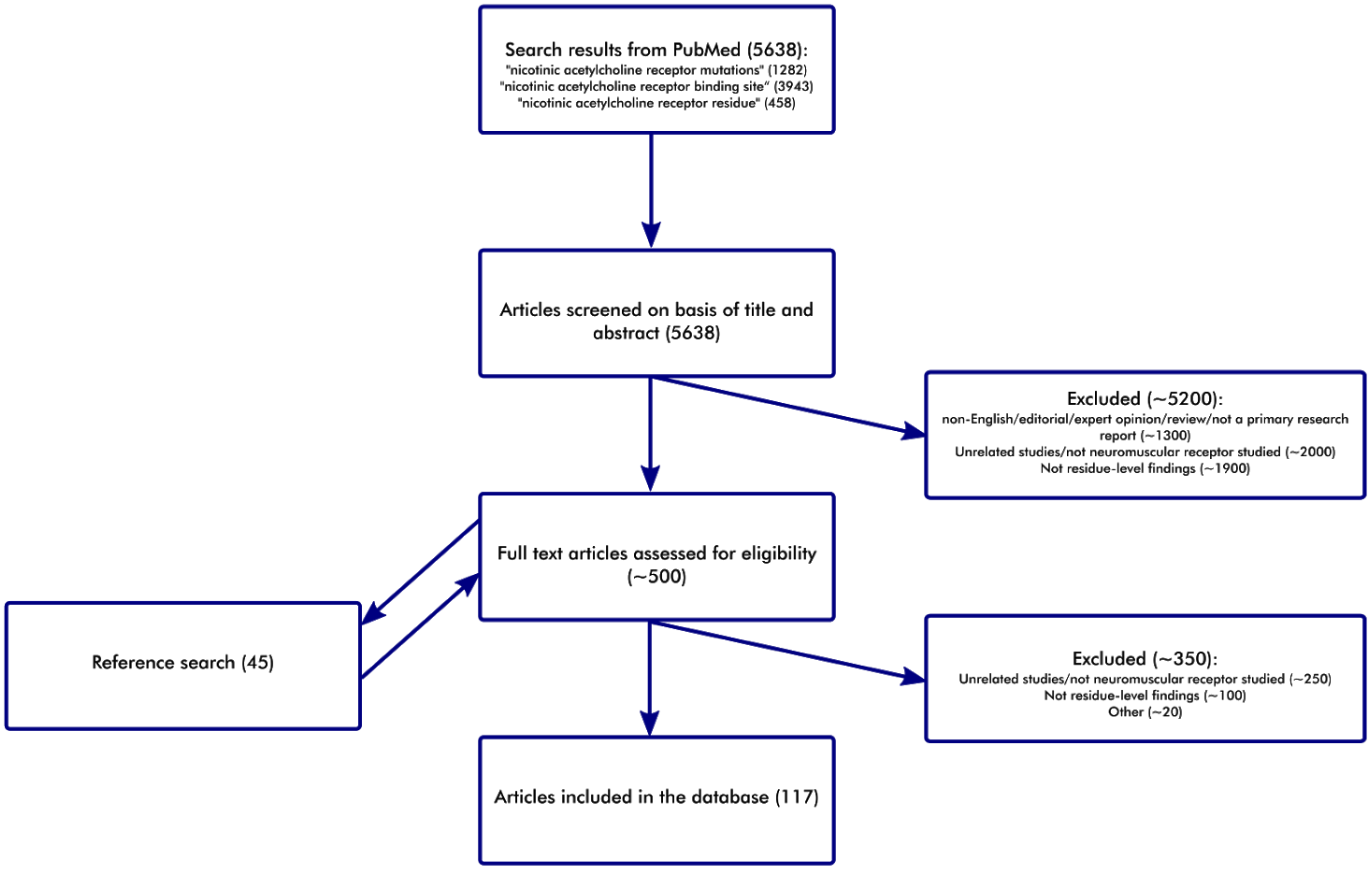


**Figure 1. Flowchart depicting the literature scanning process**

To gather a sufficient number of data points, we employed the following workflow. First, we searched PubMed for articles describing the functional role of neuromuscular nicotinic acetylcholine receptor amino acid residues. Next, we briefly reviewed the titles and abstracts and excluded a part of them based on the mentioned exclusion criteria. For the remaining articles, we reviewed the full text and excluded the additional articles based on the same exclusion criteria and on additional criterion (significant ambiguity in the full text, such as inconsistent residue numbering, or residue numbering not corresponding to the canonical sequences of nAChR subunits, or chain IDs/subunit types not specified properly etc). The data from a total of 117 articles were included in the database.

### PDB search

On 23.11.2019, we searched PDB in order to find all complete nAChR structures. First search was performed with the following search parameters:

*Text search: “nicotinic acetylcholine receptor”, Chain length: from 300, Source Organism Browser (NCBI): Eukaryota.*

Second search was performed with the following search parameters:

*Text search: “nicotinic receptor”, Chain length: from 300, Source Organism Browser (NCBI): Eukaryota.*

After removing the duplicates and manual inspection, the total of 10 PDB structures were added to the NAChRDB (PDB IDs: 6PV7, 6PV8, 6CNJ, 6CNK, 5KXI, 4BOR, 4BOT, 4AQ9, 4AQ5, 2BG9).

### UniProt search

On 23.11.2019, we searched UniProt KB in order to find the complete nAChR sequences. The following search query was used:

*taxonomy:"Bilateria [33213]" "nicotinic acetylcholine receptor" NOT toxin length:[300 TO *] family:"ligand-gated ion channel tc 1 a 9 family acetylcholine receptor tc 1 a 9 1 subfamily" keyword:"Ion channel [KW-0407]" keyword:"Ligand-gated ion channel [KW-1071]" keyword:"Receptor [KW-0675]" keyword:"Ion transport [KW-0406]" keyword:"Transport [KW-0813]"*

Our search retrieved 91 UniProt entries in total. Five of them were excluded because it was not possible to infer which subunit type, according to commonly recognized typology (alpha, beta, delta, gamma, epsilon) they correspond to (UniProt IDs: Q9U298, Q60S81, P13908, P18257, P54247). Three other UniProt entries corresponding to Torpedo marmorata nAChR (subunits delta, beta, gamma, UniProt IDs: Q6S3H8, Q6S3I0, Q6S3H9) were missing from the search since they are not classified as members of acetylcholine receptor (TC 1.A.9.1) subfamily in UniProt KB. Since they are widely recognized as subunits of Torpedo nAChR, they were manually selected and included into the NAChRDB.
After the manual inspection, the data from a total of 89 UniProt entries was included in the NAChRDB (UniProt IDs: Q6S3H8, Q6S3I0, Q6S3H9, P43144, Q9JLB5, Q9UGM1, Q9GZZ6, P25162, P17644, P04755, P09478, Q9PTS8, P17787, P32297, P43681, P12390, P09484, P36544, P30926, P04757, P12392, Q9ERK7, P09483, Q8R4G9, P30532, Q8R493, O70174, P09482, Q9N587, G5EG88, P19370, P25108, Q2MKA5, Q05901, P22770, Q04844, P04760, P26152, Q8BMN3, Q9R0W9, P02710, P25109, P43679, P02718, P25110, P09660, P49581, P20782, P49579, P02713, P45963, P09480, P02712, Q07001, P20420, Q15822, Q07263, P09481, P12389, P18845, P02714, P26153, Q15825, P43143, Q05941, P12391, P02717, P18916, P54244, P48181, P49582, Q91X60, Q93149, Q68RJ7, Q27218, O76554, Q23022,

P48182, P04756, P02708, P09690, P91766, P09479, Q9I8C7, Q8JFN7, P07510, P23414, P02716, P11230).

### Alignment

Multiple sequence alignment of all protein sequences in the database were performed using Clustal W with the default settings (Thompson et al., 1994) and manually inspected upon completion.

### Partial atomic charges calculation

The starting molecular structures of muscle nAChR were obtained from PDB entries 4AQ5 and 4AQ9 (Unwin and Fujiyoshi, 2012). TRITON program (Prokop *et al.*, 2000) was used to protonate both structures and correct the known minor structural issues. The web server elNémo (Suhre and Y.-H. Sanejouand, 2004), with default settings, was used to calculate the low frequency normal modes, and generate normal mode perturbed models of muscle nAChR. In *elNémo*, normal mode calculation is based on the harmonic approximation of the potential energy function around a minimum energy conformation, where a single-parameter Hookean potential is used. The Elastic Network Model (Tama and Sanejouand, 2001; Suhre and Y. H. Sanejouand, 2004) is then applied to generate models that are perturbed with a given amplitude in the direction of a single normal mode. In our study, a total of 11 low energy models of each of the two muscle nAChR structures available in the PDB were generated, perturbed in the direction of the 5 lowest frequency normal modes (parameters: minimum perturbation, -100; maximum perturbation, 100; step size, 20; and cutoff for the elastic interaction, 8 Å). We then chose for further analysis the two most distinct models (highest RMSD with respect to C-α atoms). The channel radius profiles were computed for these two models using the web server ChExVis (Masood *et al.*, 2015), with default settings. The model with a wider channel profile in the known gating area was designated as the *open* conformer, and the other model was designated as the *closed* conformer. The 3D models of the *closed* and *open* conformers of muscle nAChR used to derive the results presented in this study are available in .pdb format (Files S1.pdb and S2.pdb).

The protonated structures were also carefully inspected in PyMol v2.2.0.0 (Schrödinger, LLC, 2015). The atomic charges were computed using Atomic Charge Calculator, or ACC (Ionescu *et al.*, 2015), with the following setup: total charge for each conformer, -16 *e*; parameter set, EX-NPA 6-31Gd PCM (Ionescu *et al.*, 2013); computation method, Full EEM. ACC relies on the Electronegativity Equalization Method (Mortier *et al.*, 1986) to compute atomic charges which respond to changes in molecular conformation. Therefore, despite having the same total charge and using the same computation setup, the two conformers exhibited different atomic charges. The atomic charges for the *closed* and *open* conformers of muscle nAChR used to derive the results presented in this study are available in .csv format (S3.csv file). Additionally, partial atomic charges were calculated employing another EEM parameter set (Raček *et al.*, 2018), data available in S4.csv file. Additionally, partial atomic charges were computed utilizing other than EEM charge calculation method – charge equilibration (QEq) (Rappe and Goddard, 1991; Raček *et al.*, 2018, 2016), data available in S5.csv file.

In EEM ACC implementation for each amino acid residue *r*, the total charge $Q_r$ was obtained as the algebraic sum of the atomic charges *qi* on all the atoms that belong to residue *r*:

$$Q_r = \sum_{i=1}^{n_r} qi \quad (1)$$

where $n_r$ is the number of atoms in residue *r*. The significance of residue *r* was evaluated based on the absolute difference $D_r$ between its charge in the open conformer $Q_{ro}$, and its charge in the closed conformer $Q_{rc}$:

$$D_r = \left| Q_{ro} - Q_{rc} \right| \quad (2)$$

Residue *r* was considered significant if its $D_r$ lay above the upper inner fence of the distribution of absolute differences for the entire molecule:

$$D_r \geq D^{75} + 1.5 \cdot \left( D^{75} - D^{25} \right) \quad (3)$$

where $D^{75}$ and $D^{25}$ represent the upper and lower quartiles, respectively, of the distribution of $D_r$ values for all residues in muscle nAChR. This significance threshold can be further extended to:

$$D_r \geq D_{ave} + 2 \bullet \sigma \quad (4)$$

where $D_{ave}$ and $\sigma$ represent the average value and the standard deviation, respectively, of the distribution of $D_r$ values for all residues in muscle nAChR.

### Channel lining residues identification

The list of channel lining residues for each PDB entry in the NAChRDB were obtained from ChannelsDB (Pravda *et al.*, 2018) using the following API request: https://webchem.ncbr.muni.cz/API/ChannelsDB/PDB/*pdb_id*

where *pdb_id* is a PDB accession number (ID)

### References


Ionescu,C.-M. *et al.* (2015) AtomicChargeCalculator: interactive web-based calculation of atomic charges in large biomolecular complexes and drug-like molecules. *J. Cheminform.*, **7**, 50.

Ionescu,C.-M. *et al.* (2013) Rapid Calculation of Accurate Atomic Charges for Proteins via the Electronegativity Equalization Method. *J. Chem. Inf. Model.*, **53**, 2548–2558.

Masood,T. Bin *et al.* (2015) CHEXVIS: A tool for molecular channel extraction and visualization. *BMC Bioinformatics*, **16**.

Mortier,W.J. *et al.* (1986) Electronegativity-equalization method for the calculation of atomic charges in molecules. *J. Am. Chem. Soc.*, **108**, 4315–4320.

Pravda,L. *et al.* (2018) ChannelsDB: database of biomacromolecular tunnels and pores. *Nucleic Acids Res.*, **46**, D399–D405.

Prokop,M. *et al.* (2000) TRITON: In silico construction of protein mutants and prediction of their activities. *Bioinformatics*, **16**, 845–846.

Raček,T. *et al.* (2018) Empirical methods for calculation of partial atomic charges – applicability for proteins? In, *ENBIK2018*.

Raček,T. *et al.* (2016) NEEMP: software for validation, accurate calculation and fast parameterization of EEM charges. *J. Cheminform.*, **8**, 57.

Rappe,A.K. and Goddard,W.A. (1991) Charge equilibration for molecular dynamics simulations. *J. Phys. Chem.*, **95**, 3358–3363.

Schrödinger, LLC (2015) The {PyMOL} Molecular Graphics System, Version~1.8.

Suhre,K. and Sanejouand,Y.-H. (2004) ElNemo: a normal mode web server for protein movement analysis and the generation of templates for molecular replacement. *Nucleic Acids Res.*, **32**, W610-4.

Suhre,K. and Sanejouand,Y.H. (2004) ElNémo: A normal mode web server for protein movement analysis and the generation of templates for molecular replacement. *Nucleic Acids Res.*, **32**.

Tama,F. and Sanejouand,Y.H. (2001) Conformational change of proteins arising from normal mode calculations. *Protein Eng.*, **14**, 1–6.

Thompson,J.D. et al. (1994) CLUSTAL W: Improving the sensitivity of progressive multiple sequence alignment through sequence weighting, position-specific gap penalties and weight matrix choice. *Nucleic Acids Res.*, **22**, 4673–4680

Unwin,N. and Fujiyoshi,Y. (2012) Gating movement of acetylcholine receptor caught by plunge-freezing. *J. Mol. Biol.*, **422**, 617–634.
